# Optimization of culture and transfection methods for primary snake cells

**DOI:** 10.1101/2025.04.04.647168

**Authors:** Shoma Kuriyama, Keisuke Shigematsu, Seung June Kwon, Ryusei Kuwata, Yuji Atsuta

**Affiliations:** Department of Biology, Faculty of Science, Kyushu University, Fukuoka 819-0395, Japan; Faculty of Veterinary Medicine, Okayama University of Science, Ehime 794-8555, Japan

**Keywords:** primary snake cells, culture optimization, transcriptome, gene manipulation, chimeric limb bud

## Abstract

Snakes serve as important models for understanding how changes in genes and genome sequences drive vertebrate morphological evolution. However, the lack of established primary culture methods and gene delivery techniques for snake cells has hindered functional analyses of evolutionarily modified genes and genomic elements. Here, we optimized primary culture conditions and screened for efficient transfection methods using corn snake embryonic fibroblasts. Our culture optimization experiments revealed that TeSR medium, designed for stem cells, with fetal bovine serum supplementation, and incubation at 28℃ provided a suitable condition for primary snake fibroblasts. Transcriptome analysis further demonstrated that under this optimized condition, genes associated with cytoskeletal organization, extracellular matrix components, and sterol biosynthetic process were upregulated, likely promoting snake cell proliferation. Additionally, screening of various gene transfection methods identified three efficient approaches, including electroporation. These findings enhance the utility of snake cells and pave the way for functional analyses of genes and genomic elements using snake cell-based systems.

## 1 INTRODUCTION

Snakes have acquired a variety of unique traits through the process of adaptive evolution. They have lost their limbs, sterna, and external ears, while developing elongated bodies, highly flexible jaws, and a keen sense of smell through the Jacobson’s organ. Additionally, many snake species have evolved venom glands from parts of their salivary glands. These snake-specific traits arise during embryonic development and are of great interest from an evolutionary developmental biology perspective. However, the cellular and molecular mechanisms that govern these morphologies remain largely unknown. The limited progress in unraveling the mechanisms underlying snake-specific traits is likely due to two major challenges that must be overcome.

First, the primary challenge is the difficulty of utilizing snake embryos. Snakes exhibit seasonal reproduction, typically laying eggs only once or twice a year. Although artificial temperature control can be used to adjust the timing of egg laying, increasing the frequency remains nearly impossible. In addition, in species such as the corn snake (*P. guttatus*), which has a fully sequenced genome and is used for studying body pattern formation (Saenko *et al*., 2015; Tzika *et al*., 2024), and the Japanese striped snake (*E. quadrivirgata*), which has been used in developmental studies (Matsubara *et al*., 2016; Matsubara *et al*., 2017), extra-embryonic blood vessels are already well-developed and tightly attached to the eggshell membrane, even in freshly laid eggs. These blood vessels hinder *in ovo* manipulations commonly performed in avian embryonic experiments, making microsurgery and gene manipulation in snake embryos extremely challenging.

Secondly, another major challenge is the underdeveloped technology for generating snake tissue organoids, which could serve as alternative models for developing embryonic tissues. Although there has been a report on the successful culture of venom gland cells from venomous snakes and the construction of functional venom gland organoids (Post *et al*., 2020), there are no reports on the generation of organoids from other snake cell types. Moreover, the derivation of pluripotent cells, such as induced pluripotent stem cells (iPSCs), and the cultivation methods for tissue stem cells required for organoid generation have not yet been established for snake cells. The lack of systematic optimization of primary culture systems and gene delivery methods for snake cells is considered the root of this issue.

In this study, to contribute to solving the second issue, we used corn snake embryonic fibroblasts (Pg SEFs) as the main experimental model to investigate primary culture conditions and explore efficient gene manipulation methods. Regarding the culture conditions, we examined the effects of various serum concentrations and types, additives such as ovalbumin, stem cell culture medium, and temperature changes on the proliferation of snake cells. Furthermore, we conducted transcriptomic analysis to identify enriched genes and biological processes in Pg SEFs cultured under the optimized condition. Additionally, to screen for an efficient gene introduction method, we tested commercially available cationic-lipid transfection reagents, non-lipid synthetic polymer reagents, and an electroporation method. We also verified whether the optimized culture condition and gene delivery method for Pg SEFs were effective in adult corn snake fibroblasts and Japanese striped snake fibroblasts.

## 2 MATERIALS AND METHODS

### 2.1 Experimental animals

Adult corn snakes (*P. guttatus*) were acquired from a licensed reptile shop (Fukuoka, Japan). The snakes were individually housed in separate cages, fed a mouse weekly, and kept at a temperature of 20-28℃ and a humidity of 40-70%, suitable for corn snake husbandry (Pees *et al*., 2016). To obtain fertilized snake eggs, sexually mature pairs were housed together in the same cage in spring for mating. After mating, the eggs were laid approximately 45 days later and incubated at 28℃ and ∼70% humidity until they were used. Corn snake husbandry and sacrifice of a juvenile snake (7-day-old) were carried out according to the protocols approved by Kyushu University (No. Faculty of Science 1-12 and A23-315-1).

Fertilized Japanese striped snake (*E. quadrivirgata*) eggs were obtained from the Japan Snake Institute (Gunma, Japan). The eggs were kept at 28℃ and ∼70% humidity. Adult Japanese striped snakes were captured in the wild, and were humanly sacrificed following an approved protocol at Okayama University of Science.

Fertilized chicken eggs of Hypeco nera were purchased from Yamagishi (Wakayama, Japan). Chicken eggs were incubated at 38.5℃, and embryos were staged according to the Hamburger-Hamilton (HH) stages (Hamburger & Hamilton, 1951). Experiments utilizing chicken embryos were conducted under the ethical approval of Kyushu University (No. A24-070-0).

### 2.2 Culture media

Dulbecco’s Modified Eagle Medium (DMEM) and fetal bovine serum (FBS) were purchased from FUJIFILM and Biowest, respectively. FBS/DMEM used in this study contained FBS, Non-essential amino acid (NEAA, Thermo Fisher Scientific), β-mercaptoethanol (βME, Thermo Fisher Scientific), Vitamin solution (Thermo Fisher Scientific), and Penicillin-Streptomycin (Thermo Fisher Scientific). For CS/DMEM, chicken serum (Biowest) was used instead of FBS. TeSR-E6

(Veritas) was used for FBS/TeSR or CS/TeSR instead of DMEM. Ovalbumin (Sigma-Aldrich) and human insulin (FUJIFILM) were used at concentrations of 2.5% and 1.7 μM, respectively.

### 2.3 Collection of snake and chicken fibroblasts

For collecting corn and striped snake embryonic fibroblasts (Pg SEFs and Eq SEFs), embryos at 8 days post oviposition (DPO8) were harvested, and internal organs were removed. The trunk regions were dissected out and incubated in 0.25% Trypsin/EDTA (Gibco) at 37℃ for 20 min, occasionally pipetting to suspend the tissue. After quenching the enzymatic activity of trypsin by adding 15% FBS (Thermo Fisher Scientific)/DMEM (FUJIFILM), the cells were plated and cultured in 15% FBS/DMEM at 32℃ until they reached confluence. After being passaged twice, the SEFs were either used for experiments to optimize culture conditions or preserved in Bambanker (NIPPON Genetics) at -80℃ until use.

To obtain corn snake adult fibroblasts (Pg SAFs), liver tissues were dissected out from a juvenile corn snake. The tissues were incubated in 0.25% Trypsin/EDTA (Gibco) at 37℃ for 30 min with occasional pipetting. After dissociation, the cells were seeded in 10-cm dishes (IWAKI) and cultured in 15% FBS/DMEM at 28℃ until they reached confluence. After being passaged twice, the Pg SAFs were either used for experiments to optimize culture conditions or preserved in Bambanker at -80℃ until use. The medium used for culturing Pg SAFs did not contain EGF, insulin, glucagon or other components necessary for primary hepatocyte culture (Kaur *et al*., 2023). Additionally, from a morphological perspective, the remaining cells were considered to be fibroblasts.

For collecting adult striped snake fibroblasts (Eq SAFs), liver tissues were dissected out from an adult striped snake, which was caught in the wild. The tissues were incubated in 0.25% Trypsin/EDTA (Gibco) at 37℃ for 30 min with occasional pipetting. After dissociation, the cells were seeded in 10-cm dishes (IWAKI) and cultured in 15% FBS/DMEM at 28℃ until they reached confluence. After being passaged six times, the Eq SAFs were either used for experiments to optimize culture conditions or preserved in Bambanker at -80℃ until use.

Primary chicken embryonic fibroblasts were prepared as previously described (Atsuta *et al*., 2022). Briefly, posterior parts of HH32 embryos were dissected out and trypsinized. The dissociated cells were cultured in 15% FBS/DMEM and have been passaged two times. The cells have been kept in Bambanker at -80℃ until use.

All cells were passaged at a dilution rate of 1/10.

### 2.4 Transcriptomic analyses

Total RNA was extracted from SEFs cultured at 20℃, 28℃ or 37℃ for one week using NucleoSpin RNA Plus XS (TAKARA). Library preparation for BRB-Seq (Alpern *et al*., 2019), sequencing (NovaSeq 6000, Illumina), raw reads quality control and mapping were performed by Transcriptomics Kenkyukai (Fukuoka, Japan). UMI-tools (ver.1.1.4; Smith *et al*., 2017) were used to extract barcode reads. Adaptor and low-quality sequences were filtered using Trim Galore (ver.0.6.10; Krueger, 2015), and reads were aligned to the CU_Pguttatus_1 genome (GCF_029531705.1; https://www.ncbi.nlm.nih.gov/datasets/genome/GCF_029531705.1/) using HISAT2 (ver.2.2.1; Kim *et al*., 2019). Mapped reads were quantified with featureCounts (ver.2.0.6; Liao *et al*., 2013), and differential expression was analyzed using DESeq2 (ver.1.38.3; Love *et al*., 2014). Volcano plots for differentially expressed genes (|log2FC | > 1.5 and padj <0.00001) were generated by EnhancedVolcano (ver1.16.0; Blighe et al., 2018). Gene Ontology (GO) analysis was performed using the corn snake GO annotation file (gaf; https://www.ncbi.nlm.nih.gov/datasets/genome/GCF_029531705.1/) and clusterProfiler (ver.4.6.2; Yu *et al*., 2012). The “enrichGO” command with Benjamini and Hochberg’s method for p-value adjustment and p-, q-value cut-off of 0.05 was used for GO enrichment analysis. A heatmap of log2FCs compared to 28℃ condition was generated using pheatmap (ver.1.0.12; Kolde, 2018).

### 2.5 Plasmid DNA transfection and electroporation

pCAGGS-EGFP, pCAGGS-rtTA2^S^-M2 and pT2A-TRE-FGF8-EGFP were described elsewhere (Watanabe *et al*., 2007). The transfection protocols for each reagent are described below. All transfection experiments were performed using 24-well plates (Nunc), and 500 μL of Opti-MEM (Thermo Fisher Scientific) was pre-added to each well. The media containing transfection reagents were replaced with 15% FBS/DMEM, 24 hrs after the start of transfection.

Lipofectamine 2000 (Thermo Fisher Scientific); 1 μL Lipofectamine 2000 was added to 50 μL Opti-MEM and mixed by pipetting. The mixture was then combined with 50 μL Opti-MEM containing 1 μg pCAGGS-EGFP and resuspended thoroughly. After incubation at room temperature (RT) for 15 min, the mixture was added to the cells.

Lipofectamine 3000 (Thermo Fisher Scientific); 1.5 μL Lipofectamine 3000 was added to 50 μL Opti-MEM and mixed by pipetting. The mixture was combined with 50 μL Opti-MEM containing 1 μg pCAGGS-EGFP and 1 μL P3000, and resuspended thoroughly. After incubation at RT for 15 min, the mixture was added to the cells.

TransfeX (ATCC); 1 μL TransfeX and 1 μg pCAGGS-EGFP were added to 50 μL Opti-MEM. After incubation at RT for 15 min, the mixture was added to the cells.

Xfect (TAKARA); 1 μg pCAGGS-EGFP were added to 30 μL Reaction Buffer, and then 0.3 μL Xfect polymer. After incubation at RT for 10 min, the mixture was added to the cells.

FuGENE HD (Promega); 1 μL FuGENE HD was added to 50 μL Opti-MEM and mixed by pipetting. The mixture was then combined with 50 μL Opti-MEM containing 1 μg pCAGGS-EGFP and resuspended thoroughly. After incubation at RT for 15 min, the mixture was added to the cells.

Effectene (QIAGEN); 1 μg pCAGGS-EGFP and 1.6 μL Enhancer were added to 60 μL Buffer EC and mixed by pipetting. After incubation at RT for 5 min, 5 μL Effectene Reagent was added to the mixture. After further incubation at RT for 10min, the mixture was added to the cells.

Polyethylenimine Max (PEI, Polysciences); 2 μg pCAGGS-EGFP and 6 μL PEI (1 mg/mL) were added to 500 μL Opti-MEM, and the mixture was incubated at RT for 10 min, then added to the cells.

For electroporation to snake cells, ∼1 x 10^5^ cells were resuspended in 10 μL T buffer (Thermo Fisher Scientific) containing 1 μg DNA plasmids (1 μg pCAGGS-EGFP; 400 ng pCAGGS-rtTA-M2 and 600 ng pT2A-TRE-FGF8-EGFP), and electroporated using Neon transfection system (Thermo Fisher Scientific) with three pulses of 1,000 V (for SEFs) or 1,300 V (for SAFs) for 10 ms.

Observation and sorting of EGFP-expressing cells were performed 48 hrs after transfection. Images were taken using an inverted microscope IX83 (Olympus) equipped with a Zyla-4.2 Plus camera (ANDOR), and cell sorting was conducted using the SH800 (Sony).

### 2.6 Cell counting and statistical analyses

When cultured snake cells were passaged, the number of the cells was counted using a hemocytometer (AZ-ONE) or an automated cell counter (TC20, BIO-RAD). The proportion of EGFP-positive cells was measured using the SH800. Statistical analyses were performed with GraphPad Prism 10 software (GraphPad).

### 2.7 Transplantation of FGF8-expressing cells into chicken embryos

pCAGGS-rtTA2^S^-M2 and pT2A-TRE-FGF8-EGFP were introduced into DF-1 or SEF cells by electroporation. 24 hrs after doxycycline administration (1 μg/mL; FUJIFILM), cell aggregates were prepared by a hanging drop method (Rasouli *et al*., 2025). Transplantation of cell aggregates into frank regions of HH10 embryos was performed using a tungsten needle. The embryos grafted with FGF8-expressing cells were incubated at 37℃ until HH25.

## 3 RESULTS

### 3.1 Increasing fetal bovine serum (FBS) or chicken serum (CS), as well as the addition of ovalbumin, does not promote the proliferation of corn snake embryonic fibroblasts (Pg SEFs)

In this study, we investigated the optimal medium components and temperature for the primary culture of snake cells. For optimization, we used fibroblasts (Pg SEFs) derived from corn snake embryos at 8 days post-oviposition, in which tissue differentiation had progressed (Fig. 1A). The corn snake embryos are scarce (as they lay ∼10 eggs only once a year), and the number of obtainable cells is limited. Therefore, we aimed to optimize culture conditions that enable active proliferation even at low cell densities. For this reason, the initial cell number was set at 10,000 cells (Fig. 1A). We first optimized the medium components while keeping the culture temperature fixed at 32℃, which has been used for cultivating cells derived from several snake species including corn snakes (Tillis *et al*., 2023).

**FIGURE 1.**
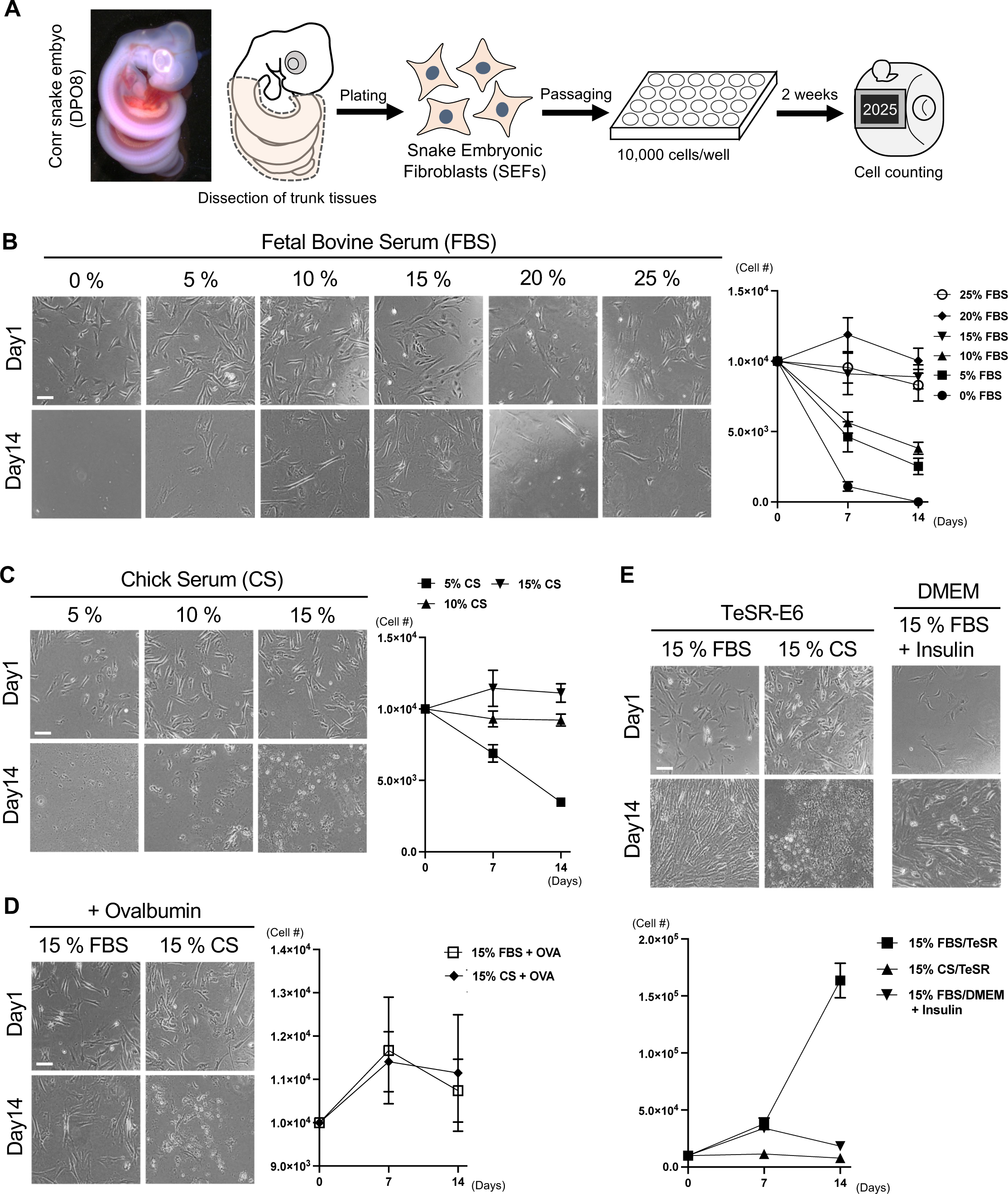
Optimization of medium composition that stimulates proliferation of corn snake embryonic fibroblasts (Pg SEFs) (A) The trunk region of the corn snake embryo at DPO8 (8 days post oviposition) was dissected, and fibroblasts (snake embryonic fibroblasts: SEFs) were harvested. After the second passage, the cells were seeded at an initial density of 10,000 cells/well and cultured in each condition for two weeks. The cell count was then measured. (B-D) DMEM was used as the basal medium. (B) The effect of fetal bovine serum (FBS) concentration on the growth of SEFs. (C) The effect of chicken serum (CS) concentration on the growth of SEFs. (D) The addition of ovalbumin fails to promote the proliferation of SEFs. (E) The effects of switching the basal medium from DMEM to TeSR-E6 and the addition of insulin. The sample size (n) for each culture condition was 6. Statistical results with One-way ANOVA for (B), (C), and (E) are shown in Fig. S2. Scale bars: 100 μm.

We tested the effect of different concentrations of FBS, which contains proteins, vitamins, and lipids beneficial for cell culture, with DMEM, a basal medium commonly used for mammalian and avian cell culture. Chicken embryonic fibroblasts (CEFs) used as positive controls showed a significant increase in cell number when cultured in 10% and 15% FBS/DMEM (Fig. S1A). When Pg SEFs were cultured for two weeks in an FBS-free medium, all cells did not survive (Fig. 1B). This indicates that, similar to other cell types, serum-derived nutrients are essential for Pg SEF survival. Using FBS concentration above 15% maintained the cell number; however, did not result in significant proliferation (Fig. 1B; Fig. S2A). We also examined whether the proliferation of Pg SEFs could be stimulated by taking advantage of CS and ovalbumin (Fig. 1C, D). We observed no significant increase from the initial cell number, despite a high concentration of CS (15%) (Fig. 1C; Fig. S2B). In the presence of CS, numerous lysosome-like vesicles were observed, suggesting that an immune response or cellular damage may have occurred, potentially inhibiting cell proliferation. Similarly, the addition of albumin demonstrated no efficacy for Pg SEF culture (Fig. 1D).

### 3.2 The TeSR-E6 medium in combination with FBS boosts the proliferation of Pg SEFs

Next, the basal medium was switched from DMEM to TeSR medium, which contains beneficial supplements such as F-12 (Ham, 1965), antioxidants (selenium and transferrin), and is used for stem cell culture (Ludwig *et al*., 2006). Typically, serum is not required in TeSR medium for stem cell culture. However, in this case, we used 15% FBS or CS to promote SEF proliferation more effectively. As a result, similar to its use with DMEM, 15% CS/TeSR medium did not promote the proliferation of Pg SEF (Fig. 1E; Fig. S2C). However, the combination of 15% FBS and TeSR significantly increased the cell number by approximately 16-fold (Fig. 1E; Fig. S2C).

To investigate which factor in TeSR contributes to this growth-enhancing effect, we focused on insulin, which is known to stimulate fibroblast proliferation (Monaco *et al*., 2009). When insulin was added to 15% FBS/DMEM, the cell number increased, although not to the same extent as when using TeSR (Fig. 1E; Fig. S2C). This suggests that insulin plays a role partially in the growth-enhancing effect of TeSR on Pg SEFs. Based on these results, we decided to use 15% FBS/TeSR medium for culturing snake cells.

### 3.3 Culturing at 28 ℃ is suitable for Pg SEF expansion

Following this, we explored the optimal temperature for Pg SEF culture. 28℃ is generally considered suitable for corn snake husbandry, and the snakes remain moderately active even at 20℃. Additionally, 37℃ is commonly used for expanding mammalian and avian cells. Thus, we tested five temperature conditions (20℃, 24℃, 28℃, 32℃, and 37℃) to determine the optimal temperature for culturing Pg SEFs (Fig. 2A). At 37℃, the cells survived; however, no significant increase, as seen in CEFs, was observed (Fig. 2A; Fig. S1B; Fig. S2D). Of interest, even at 20℃, when all CEFs died after two weeks, the cell number of Pg SEFs was maintained (Fig. 2A). In addition to 32℃ mentioned above, cell proliferation was also promoted at 24℃ and 28℃ (Fig. 2A; Fig. S2D). Notably, among tested temperatures, cell proliferation of Pg SEFs was most promoted at 28℃, reaching approximately 50-fold increased cell number (Fig. 2A). Along with the results obtained using 15% FBS/TeSR, Pg SEFs also proliferated at 28℃ with 15% FBS/DMEM (Fig. 2A), indicating that 28℃ is the optimal temperature for Pg SEF culture.

**FIGURE 2.**
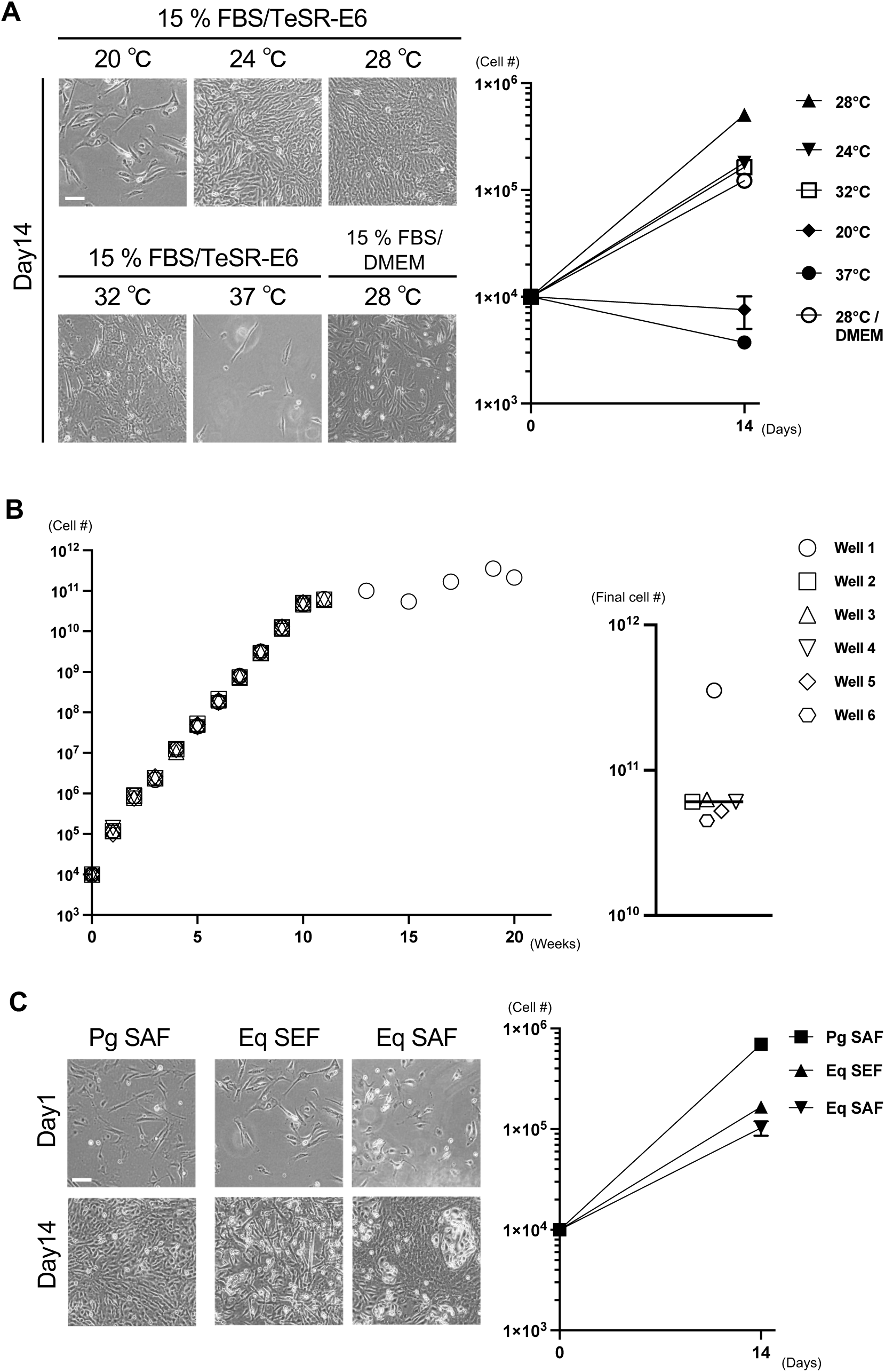
Culture at 28℃ enables the stable expansion of snake cells. (A) The corn snake SEFs (Pg SEFs) were cultured under various temperature conditions using 15% FBS/TeSR-E6 or 15% FBS/DMEM as the culture medium (n = 6 each). (B) The Pg SEFs were cultured in 15% FBS/TeSR-E6, passaged weekly, and maintained until they ceased dividing and died. The plot on the left indicates the changes in cell number over time. The right plot displays the estimated final cell number for each sample. (C) The adult corn snake fibroblasts (Pg SAFs), the embryonic and adult Japanese striped snake fibroblasts (Eq SEFs and Eq SAFs) were cultured in 15% FBS/TeSR-E6 for two weeks (n = 6 each). Statistical results with One-way ANOVA for and (C) are shown in Fig. S2. Scale bars: 100 μm.

We further tested whether the defined culture condition enables a long-term expansion of Pg SEFs. As shown in Figure 2B, Pg SEFs cultured under the defined condition showed linear growth up to 10 weeks and haves been maintained for 10-20 weeks (mean: 12.17 weeks, 11.5 passages). The final cell number was successfully increased from 1 x 10^4^ to ∼6.1 x 10^10^ (Fig. 2B), demonstrating a robust proliferative capacity. Therefore, the culture condition identified in this study were shown to be adequate for the long-term culture and large-scale expansion of Pg SEFs.

### 3.4 The defined culture condition is also optimal for adult corn snake and Japanese striped snake cells

As the next step, we investigated if the defined condition is useful for culturing cells from an adult corn snake and the Japanese striped snakes (*E. quadrivirgata*) that belong to the same Colubridae family as the corn snake. The striped snake has been used in evolutionary developmental biological research in previous studies (Matsubara *et al*., 2014; Matsubara *et al*., 2017). Thus, if the established cell culture method is also effective for the striped snake cells, it is expected to contribute to these studies as well. Fibroblasts from the adult corn snake (Pg SAFs) showed even greater proliferation (∼70-fold) than Pg SEFs under the conditions of 15% FBS/TeSR and 28℃ (Fig. 2C). Embryonic and adult striped snake fibroblasts (Eq SEFs and Eq SAFs) showed significant increases of 16.6-fold and 10.3-fold, respectively, although not as much as the corn snake cells (Fig. 2C; Fig. S2E). These results suggest that the defined condition is widely effective for expanding cells from the Colubridae family.

### 3.5 Transcriptional dynamics due to temperature differences

To visualize global gene expression changes depending on culture temperature, we carried out RNA-Seq analysis (BRB-Seq; Alpern *et al*., 2019) with Pg SEFs cultured at 20℃, 28℃ or 37℃ (Fig. 3A). The principal-component analysis (PCA) plot of RNA-seq data revealed a clear separation among the three groups, indicating distinct transcriptomic profiles (Fig. 3B). By analyzing differentially expressed genes, we found that genes related to cytoskeleton (*Krt8*, *Krt18*, *Acta2*, *Actg2*, *Actc1*) and extracellular matrix (ECM) (*Col1a1*, *Col1a2*, *Col3a1*, *Col5a1*, *Col5a2*, *Col7a1*) were highly upregulated at 28℃, compared to 20℃ and 37℃ conditions (Fig. 3C, E; Fig. S3; Fig. S4). On the other hand, *Mrnip*, whose gene product is required for DNA double-strand break repair (Wang *et al*., 2022), was upregulated both at 20℃ and 37℃ (Fig. 3C; Fig. S3; Fig. S4), suggesting that the cells cultured at these temperatures harbored more DNA damage than those at 28℃.

**FIGURE 3.**
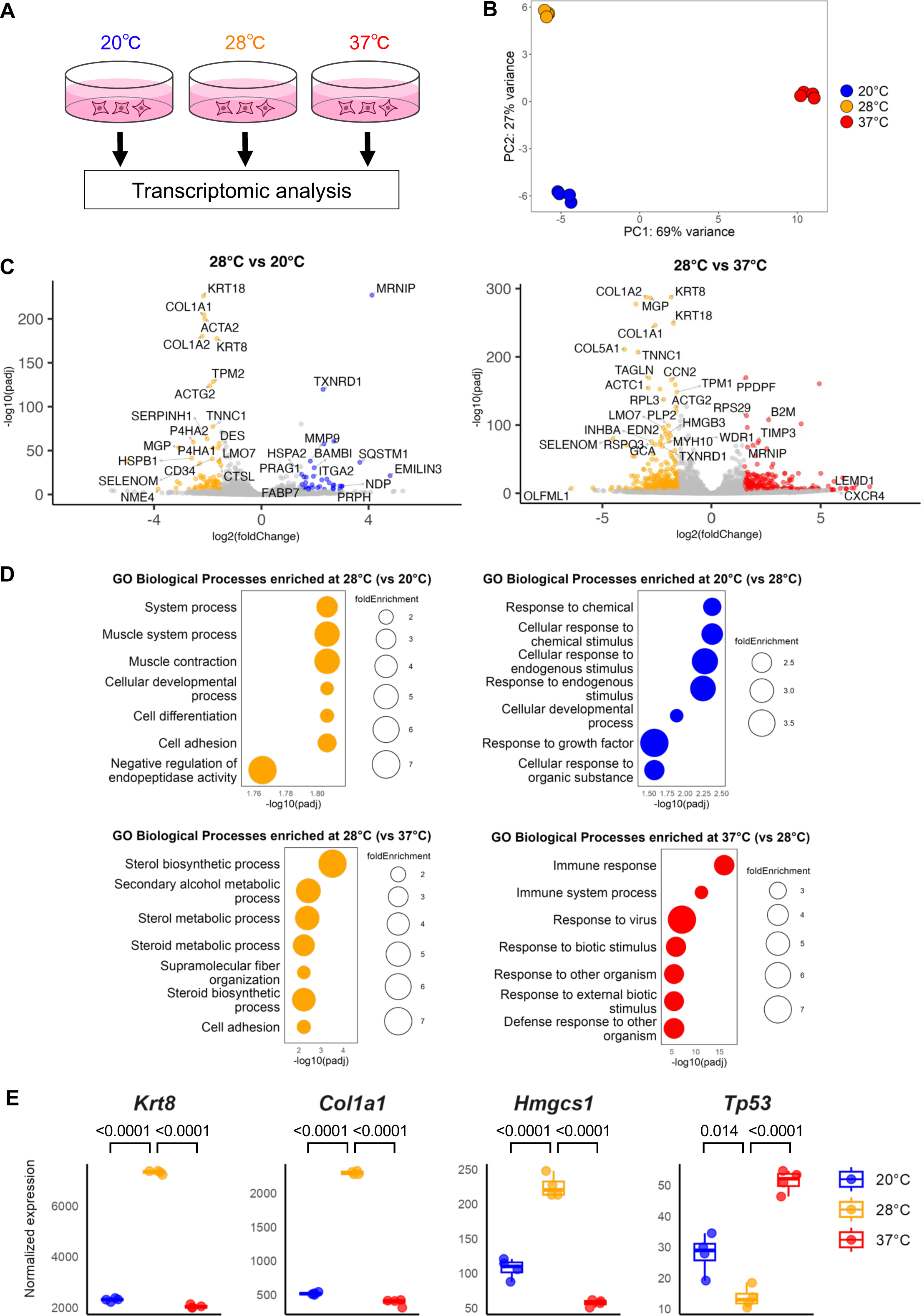
Transcriptomic analysis reveals upregulated cellular development and metabolism at 28℃. (A) Experimental design. The SEFs were cultured at 20℃, 28℃ or 37℃ for one week before being subjected to RNA-Seq analysis. (B) PCA analysis showing clear separation between temperature conditions (blue: 20℃, orange: 28℃, red: 37℃). (C) Volcano plot comparing 28℃ vs 20℃ (left) and 28℃ vs 37℃ (right). Genes with values under threshold are colored gray, while genes exceeding the threshold are colored based on the temperature condition. See also Table S1 and S2 for additional differentially expressed genes. (D) GO Biological Process terms most enriched at each temperature condition. See also Table S3 for additional enriched GO terms. (E) Box plots showing the differential expression of *Krt8*, *Col1a1*, *Hmgcs1* and *Tp53*. Normalized expression and adjusted p-values are derived from DESeq2.

To describe and compare the significant biological process in cultured Pg SEFs, we performed Gene Ontology (GO) analysis taking advantage of the corn snake GO annotation file (Fig. 3D). GO biological processes related to intracellular organization, cytoskeletal system, morphology and adhesion were enriched at 28℃ compared to 20℃ (Fig. 3D). In addition to cytoskeleton and adhesion, GO biological processes involved in sterol synthesis and alcohol metabolism were significantly enriched at 28℃, relative to the 37℃ condition (Fig. 3D). Among the genes constituting these biological processes, the expression levels of *Hmgcs1* (3-hydroxy-3-methylglutaryl-CoA synthase 1), *Hmgcr* (3-hydroxy-3-methylglutaryl-CoA reductase), *Srebf1* (Sterol regulatory element-binding transcription factor 1), *Srebf2*, *Fdft1* (Farnesyl-diphosphate farnesyltransferase 1), *Aldh2* (Aldehyde dehydrogenase 2 family member) were particularly high at 28℃ (Fig. 3E; Fig. S4). In contrast, intriguingly, under the conditions of 20℃ and 37℃, the biological process signatures related to response to chemicals and antigens were elevated compared to 28℃ (Fig. 3D). A gene encoding the chemokine receptor *Cxcr4* was highly transcribed at 20℃ and 37℃, whereas *Cxcr5*, *Cxcl11*, and *Kitlg* were upregulated at 37℃ (Fig. 3C; Fig. S4; Table S2). Their gene products are known to be involved in inflammatory responses (Hughes & Nibbs, 2018; Rönnstrand, 2004). Corresponding to the upregulation of these immune response-related genes, an increase in *Tp53* expression and a decrease in *Mdm2* were observed at 20℃ and 37℃ (Fig. 3E; Fig. S4). Tp53, a well-known tumor suppressor, acts as a transcription factor that activates genes related to DNA repair and cell cycle arrest, and its expression is induced in response to DNA damage (Nag *et al*., 2013). Mdm2 is a negative regulator of Tp53 and is downregulated upon DNA damage (Nag *et al*., 2013). Given that *Mrnip* expression was elevated at 20℃ and 37℃, DNA replication stress, which indirectly suppresses cell proliferation, may be enhanced under low- and high-temperature conditions.

We further found that the expression of the insulin receptor (*Insr*) was upregulated at 28℃, suggesting that insulin sensitivity is potentially higher under the condition (Fig. S4). Taken together, our transcriptome analysis suggested that SEFs cultured at 28℃ exhibited active cytoskeletal reorganization and ECM secretion, both of which support cell division, while maintaining lower levels of DNA damage. Additionally, efficient uptake of nutrients such as insulin may have contributed to the enhanced cell proliferation observed under this condition.

### 3.6 Screening of efficient transfection methods/reagents for snake cells

With the culture conditions established, we next explored adequate gene delivery reagents/methods for snake cells. Using Pg SEFs, we tested seven commercially available reagents (Lipofectamine 2000, Lipofectamine 3000, TransfeX, FuGENE HD, PEI, Effectene, Xfect) and the electroporation technique (Fig. 4). An EGFP-expressing plasmid was introduced, and the percentage of EGFP-positive cells was used as the transfection efficiency (Fig. 4A; Fig. S5A). As a result, relatively high transfection efficiencies were yielded with Xfect and Lipofectamine 3000 (47.3% and 57.9%, respectively), whereas TransfeX and FuGENE exhibited poor transfection efficiencies (0.1% and 0.04%, respectively; Fig. 4A). Lipofectamine 2000, PEI, and Effectene showed higher efficiencies than TransfeX and FuGENE, yet the percentage of EGFP-positive SEFs was still limited to around 10-30% (9.7%, 19.3%, and 28.2%, respectively; Fig. 4A). Among the tested methods, an electroporation demonstrated the highest plasmid transfection efficiency (72.2%; Fig. 4A). Subsequently, we investigated whether efficient transfection could be achieved in other types of snake cells using electroporation. An EGFP-expressing vector was introduced into Pg SAFs, Eq SEFs, and Eq SAFs by electroporation, and as with Pg SEFs, the percentage of EGFP-positive cells was considered the efficiency. Electroporation to Pg SAFs and Eq SEFs resulted in efficient transfection (72.6% and 69.3%, respectively), which is comparable to Pg SEFs (Fig. 4A, B; Fig. S5B). In Eq SAFs, possibly due to the higher passage number and its lower proliferative capability among the four cell types tested, the transfection efficiency was lower (Fig .4B). However, 20.4% of EGFP-positive cells were still detected (Fig. 4B).

**FIGURE 4.**
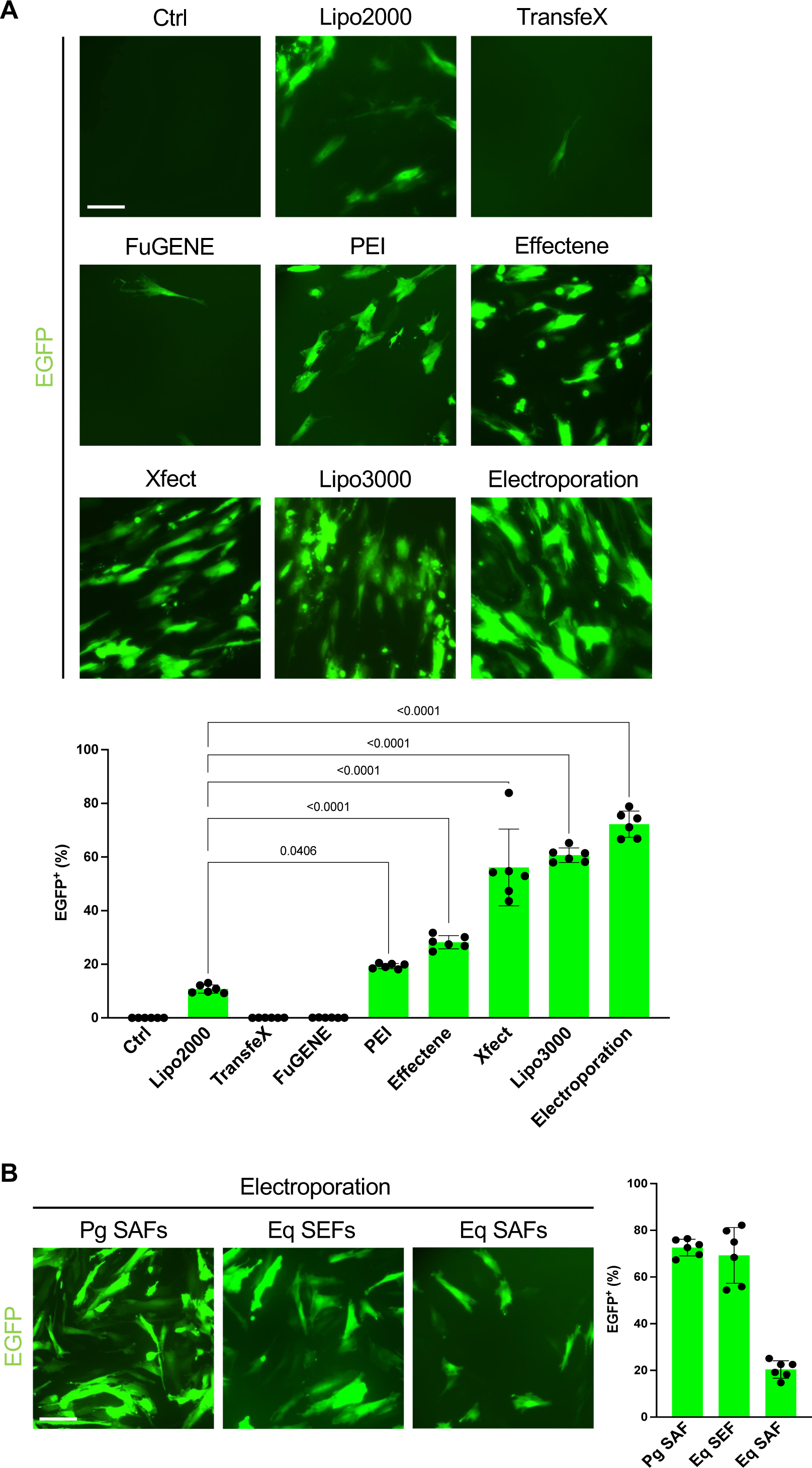
Comparison of transfection efficiencies in snake cells using various methods. (A) Images indicate Pg SEFs transfected with pCAGGS-EGFP using Lipofectamine 2000, Transfex, FuGENE HD, PEI-MAX, Effectene, Xfect, Lipofectamine 3000, or an electroporation technique. The plot shows the percentage of EGFP-positive cells measured by FACS (n = 6 each). (B) The EGFP-expressing plasmid was electroporated into Pg SAFs, Eq SEFs, or Eq SAFs. The plot represents the percentage of EGFP-positive cells measured by FACS (n =6 each). The numbers in (A) show *P* values obtained using One-way ANOVA, comparing with Lipofectamine 2000. Scale bars: 100 μm.

To summarize, we determined that two of the seven tested transfection reagents, along with electroporation, enabled efficient gene introduction into snake fibroblasts.

### 3.7 The gene of heterologous origin introduced into Pg SEFs exhibits normal function

Lastly, we examined whether an introduced transgene functions properly in snake cells. EGFP proteins appeared to localize in the cytoplasm without aggregation and exhibited normal green fluorescence (Fig. 4). Here, we further investigated a secreted factor. FGF8, a member of the FGF family, is secreted from a region called the apical ectodermal ridge (AER) during limb bud formation in tetrapods, and regulates limb growth and patterning (Lewandoski *et al*., 2000). Due to this limb-forming ability, it is known that transplantation of a bead soaked with FGF8 into the flank of a chicken embryo can induce ectopic limb formation (Crossley *et al*., 1996). Additionally, a previous study showed that python cells transplanted into a chicken limb bud could engraft without being eliminated, even in interspecies transplantation (Cohn & Tickle, 1999). Thus, by grafting an aggregate of FGF8-expressing SEFs into the lateral flank of a chicken embryo, we could determine whether SEFs properly secrete functional FGF8 based on the presence or absence of ectopic limb bud formation (Fig. 5A).

**FIGURE 5.**
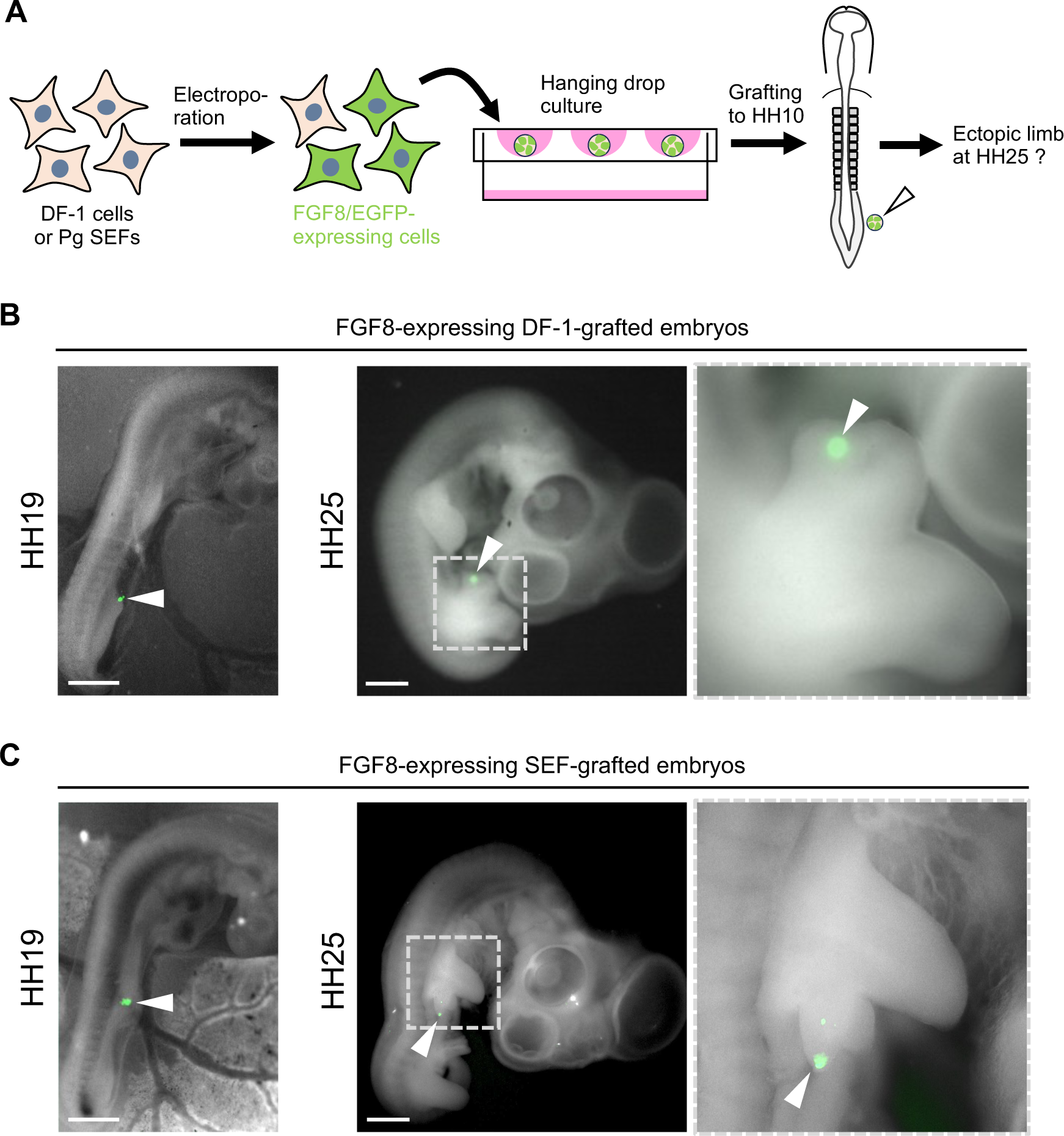
The exogenous chicken *FGF8* gene introduced into SEFs functions correctly. (A) Schematic of experiments for transplantation of the FGF8/EGFP-expressing DF-1 cells or Pg SEFs into HH10 chicken embryos. The electroporated cells were aggregated using the hanging drop culture method, and grafted into the lateral plate mesoderm of the presumptive flank region. (B) Chicken embryos at HH19 and HH25 transplanted with FGF8-expressing DF-1 cells (n = 2 for each stage). White arrowheads point the transplanted cells. (C) Chicken embryos at HH19 and HH25 transplanted with FGF8-expressing Pg SEFs (n = 3 for each stage). White arrowheads indicate the transplanted cells. Scale bars: 1 mm.

For a positive control experiment, we transfected a chicken FGF8/EGFP-expressing plasmid into chicken fibroblasts (DF-1 cells) and grafted these cells into the flank region of HH10 embryos. As expected, the grafted FGF8-DF1 cell aggregate induced an ectopic limb bud at HH25, indicating that this transplantation experiment allowed us to assess the functionality of FGF8 (Fig. 5B). The FGF8/EGFP-expressing plasmid was introduced into Pg SEFs by electroporation, and as the DF-1 experiment, the FGF8-SEF were grafted into the flank region of HH10 embryos. The grafted FGF8-SEFs induced ectopic limb outgrowth, and the grafted cells were located at the distal region of the ectopic limb bud, analogous to the AER in the endogenous limb bud (Fig. 5C). These results demonstrated that the exogenous *FGF8* gene introduced into SEFs was properly transcribed, translated, secreted, and functionally active.

## 4 DISCUSSION

In this work, through various condition optimizations, we have identified that the combination of the stem cell culture medium TeSR and FBS is suitable for the primary culture of snake fibroblasts and that culturing at 28℃ is optimal. The gene expression profiling suggested that under this culture condition, cell proliferation is promoted through the activation of cytoskeletal and ECM-related gene expression, as well as sterol biosynthesis, which is essential for mitosis progression (Fernández *et al*., 2004). In addition, we found three efficient gene delivery reagents/methods for primary cultured snake cells, and demonstrated that a heterologous transgene (chicken *FGF8*) transfected into snake fibroblasts was correctly expressed and functioned properly.

These findings would provide a foundation for deriving snake iPSCs by the overexpression of reprogramming factors, and for developing organoid techniques utilizing snake iPSCs. Furthermore, this study showed that snake cells are readily amenable to genetic manipulation. Moving forward, the use of genome editing will likely enable the direct investigation of genes involved in snake-specific morphogenesis within snake cells. Therefore, the methodology established in this study would enhance the utility of snake cells and may lead to significant progress in evolutionary developmental biology research in the future.

### 4.1 Culture conditions and temperature-responsive differentially expressed genes

This study revealed that FBS and TeSR are more effective than CS and DMEM, respectively, in promoting the proliferation of snake fibroblasts seeded at a low density. The cause of the difference in proliferation rates due to the serum type is unclear. When cultured with CS, however, numerous lysosome- or autophagosome-like vesicles were observed within the cells. This suggests that some component in CS may be triggering immune responses or cell damage.

The effectiveness of TeSR is likely, at least in part, attributed to insulin. This is supported by the finding that insulin supplementation significantly increased cell numbers even in 15% FBS/DMEM. Intriguingly, at 28℃, where cell numbers peaked, *Insr* expression was higher, implying a link between snake fibroblast proliferation and insulin signaling. However, administration of insulin alone in DMEM did not fully replicate the efficacy of TeSR, suggesting that other components, such as F-12, may also be important.

Among the temperatures tested, 28℃ was found to be optimal for primary snake fibroblasts, despite being unsuitable for CEFs. A lizard immortalized cell line was maintained at 29℃ (Samudra *et al*., 2024), and 30℃ was considered suitable for embryonic culture of Japanese striped snake eggs (Matsubara *et al*., 2014), suggesting that similar temperatures are appropriate for squamate cells. As observed in mammalian fibroblasts (Madelaire *et al*., 2022), the optimal temperature for an organism appears to be reflected in its cell culture conditions. Moreover, Pg SEFs exhibited some degree of proliferation even at suboptimal temperatures, suggesting a broader temperature tolerance, which is likely a characteristic of ectothermic animals. For instance, while the proliferation rate of CEFs dropped from approximately 507-fold to 1.9-fold (a nearly 267-fold reduction) when the culture temperature was lowered from 37℃ to 32℃, Pg SEFs showed only a 2.8-fold decrease, from 51-fold to 18-fold, when the temperature was reduced from 28℃ to 24℃.

Our transcriptome analysis revealed that the expression levels of cytoskeletal genes such as *Krt8* and *Krt18*, as well as ECM genes including *Col1a1* and *Col1a2*, were elevated at 28℃. It is known that the expression of Krt8 and Krt18 is prominent in proliferative blastema cells and malignant ovarian tumors (Corcoran & Ferretti, 1997; Barak et al., 2004), and that collagens activate fibroblast proliferation through its interaction with integrins (Pozzi *et al*., 1998). Thus, our results appear to be consistent with these findings. We also found that genes regarding the sterol biosynthetic process such as *Hmgcs1* and *Hmgcr* were enriched in SEF cultured at 28℃ compared to 37℃. In cultured rat fibroblasts, cholesterol content increases in S phase, and that is required for cell cycle progression (Singh *et al*., 2013). The activation of the sterol biosynthesis pathway, combined with the increased expression of cytokeratins and collagens, suggests that at 28℃, Pg SEFs are able to maintain an active proliferative state. Indeed, under the culture condition of 15% FBS/TeSR at 28℃, Pg SEFs could be passaged approximately 12 times. It has been reported that non-immortalized primary human dermal fibroblasts undergo a sharp increase in lysosomal enzyme characteristics and subsequently die with elevated expression of senescent markers after 12-14 passages (Cristofalo & Pignolo, 1993). The passage number of Pg SEFs is analogous to this finding.

### 4.2 Efficient gene delivery methods and their utility

We tested seven transfection reagents and electroporation, and found that Xfect (47.3%), Lipofectamine 3000 (57.9%), and electroporation (72.2%) were effective for gene delivery into Pg SEFs. Primary cells are generally more difficult to transfect compared to immortalized or cancer cell lines. In the case of MEFs, for example, reports indicated transfection efficiencies of 3-8% with Lipofectamine 2000 and FuGENE (Tsuchiya *et al*., 2016), and 12-22% with Transfectin and Lipofectamine LTX (Lee *et al*., 2017). Moreover, since SEFs appeared to have a slow cell cycle progression, which typically hampers transfection efficiency, it was unexpected that Xfect and Lipofectamine 3000 achieved efficient transfection. Additionally, similar to primary fibroblasts from chickens and humans (Atsuta *et al*., 2022; Kucharski *et al*., 2021), electroporation was the efficient gene delivery method for SEFs. To further improve gene introduction efficiency, it would be beneficial to test consecutive transfection or electroporation.

To assess whether the exogenous gene introduced was properly expressed and secreted, FGF8-expressing SEFs were grafted into chicken embryos, and the formation of ectopic limb buds was used as an indicator. The transplanted snake cells survived and engrafted without visible issues in the chicken embryos. Since it is not feasible to transplant genetically modified snake cells into snake embryos and investigate the function of genes of interest *in vivo* situations, the snake-chicken interspecies transplantation may alternatively provide a suitable *in vivo* assay system.

## AUTHOR CONTRIBUTIONS

**Shoma Kuriyama**: investigation; methodology; formal analysis. **Keisuke Shigematsu**: investigation; methodology; formal analysis. **Seung June Kwon**: investigation; methodology; formal analysis. **Ryusei Kuwata**: investigation; methodology. **Yuji Atsuta**: study design; investigation; methodology; formal analysis; writing-original draft; funding acquisition. All authors reviewed and approved the manuscript.

## ACKNOWLEDGEMENTS

We thank the Center for Advanced Instrumental and Educational Support of the Faculty of Agriculture (Kyushu University) and Transcriptomics Kenkyukai (General incorporated association, Fukuoka) for the use of SH800 and BRB-Seq, respectively. We also acknowledge Drs. Eiji Nitasaka (Kyushu University) and Daisuke Saito (Kyushu University) for helpful discussion. This work was supported by the JST FOREST Program, Grant Number JPMJFR214G (to Y.A.), JSPS KAKENHI Grant Number JP21K06201 (to Y.A.), Kato Memorial Bioscience Foundation (to Y.A.), The Naito Foundation (to Y.A.).

## DATA AND CODE AVAILABILITY

All data generated during this study are available from the corresponding author upon request. All codes used for transcriptomic analyses have been deposited in the following GitHub repository: https://github.com/seungjunekwon/Snake_culture_RNA-Seq2025-04

---

**Supplemental Figure 1.**
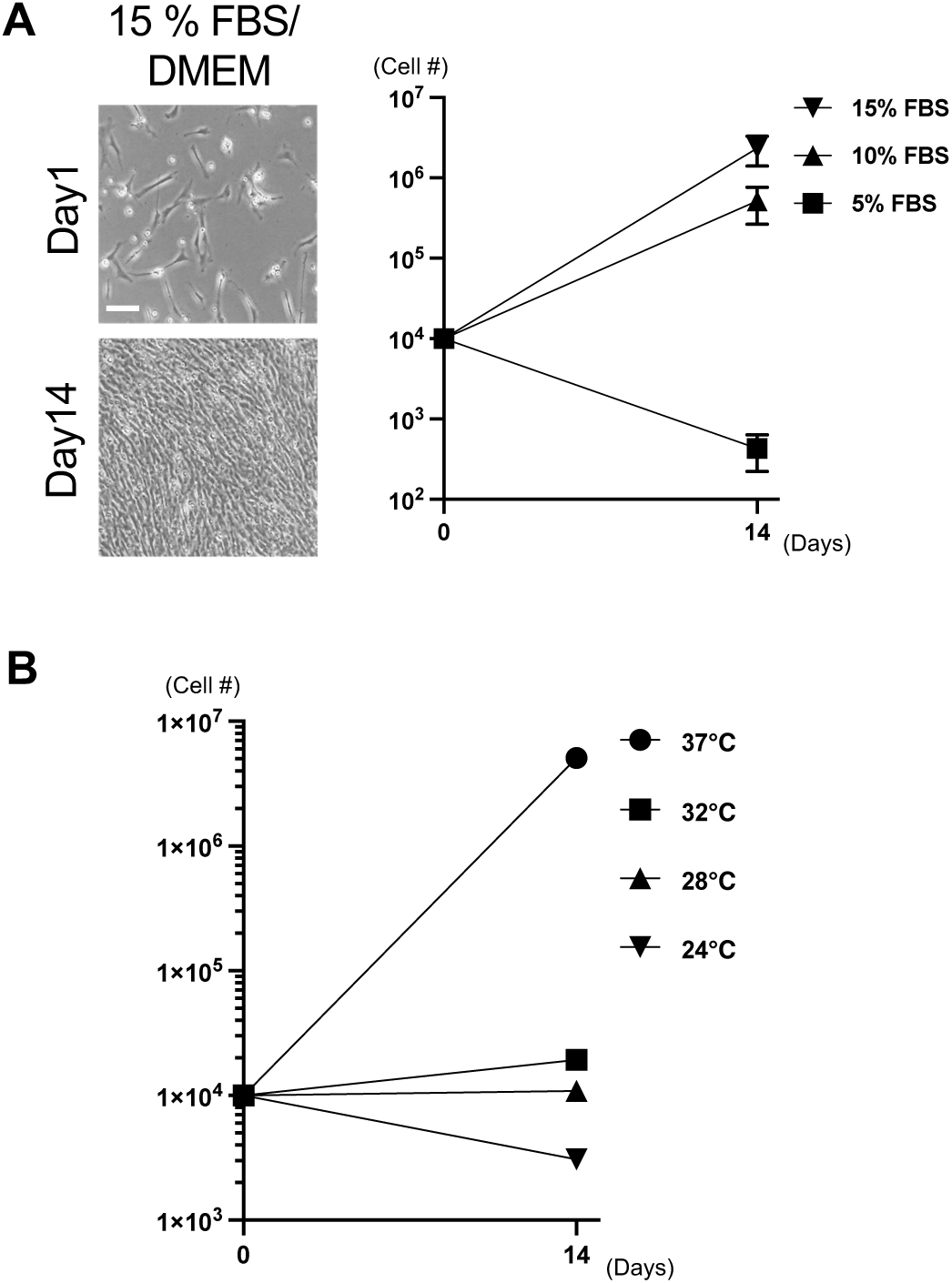
Effects of FBS concentration and temperature on the growth of chicken embryonic fibroblasts (CEFs) (A) The effect of fetal bovine serum (FBS) concentration on the growth of CEFs (n = 6 each). (B) The CEFs were cultured in 15% FBS/DMEM and under various temperature conditions for two weeks (n = 6 each). Scale bar: 100 μm.

**Supplemental Figure 2.**
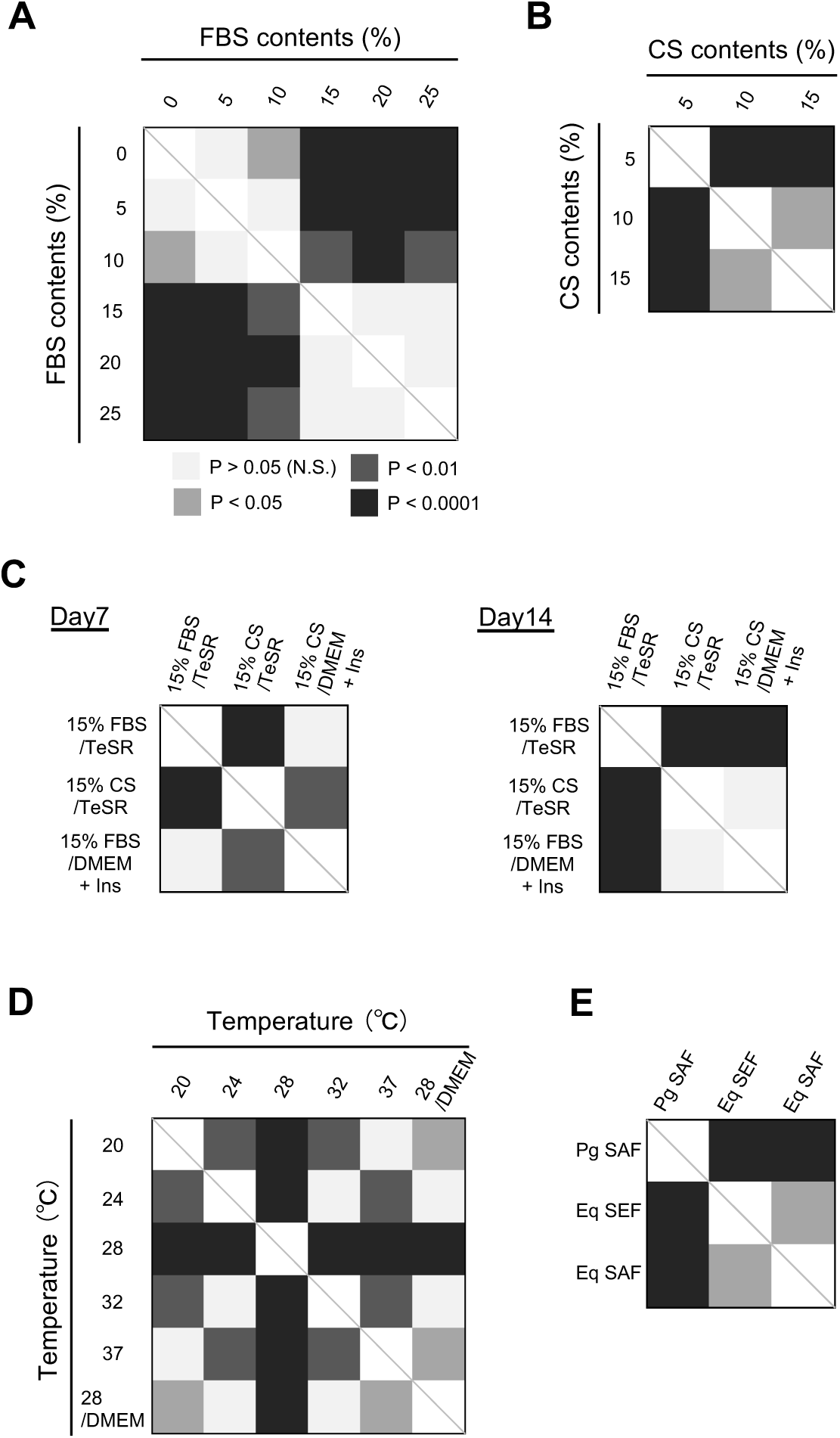
Statistical results for Figure 1 and 2. (A-E) Results corresponding to Fig. 1B (A), 1C (B), 1E (C), 2A (D), and 2C (E). All of *P* values were obtained using One-way ANOVA.

**Supplemental Figure 3.**
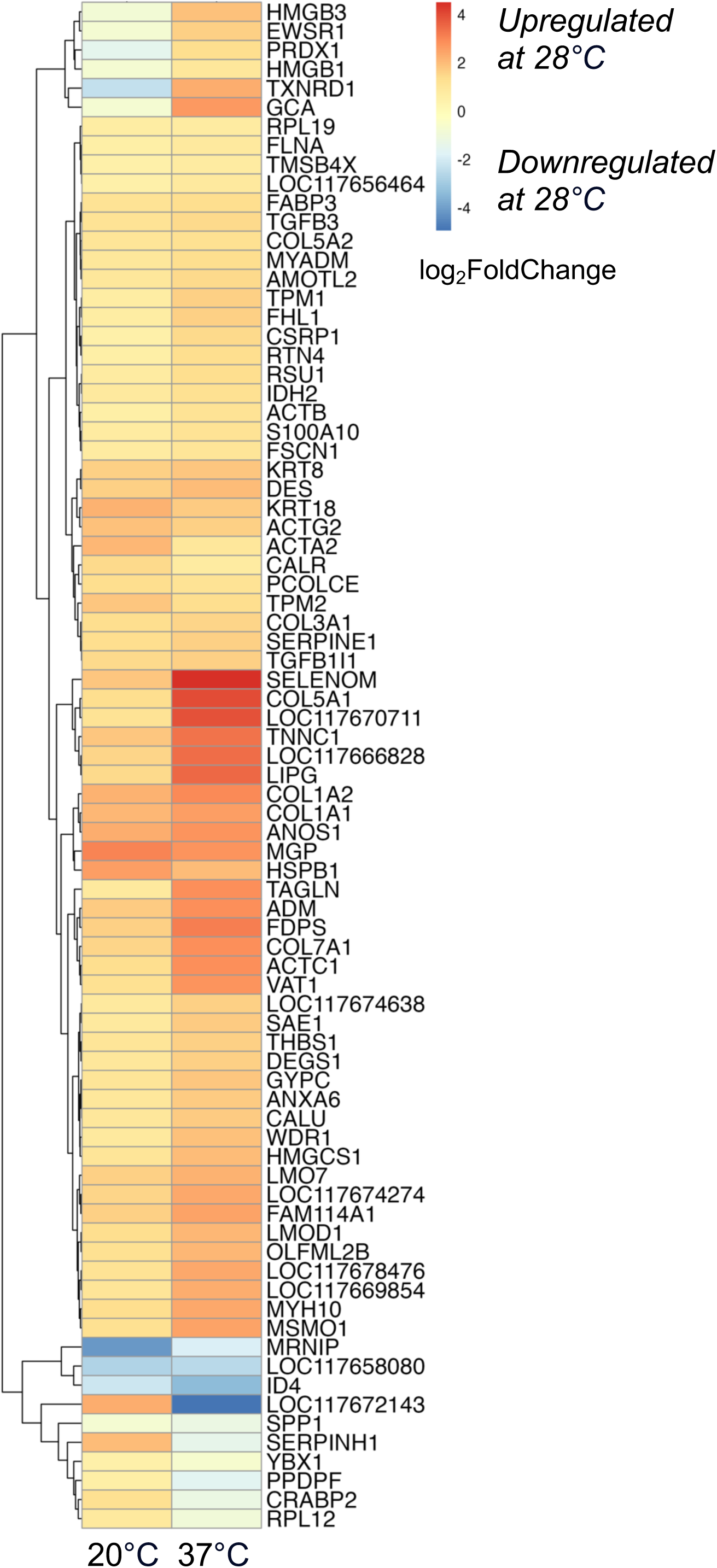
Heatmap of differentially expressed genes between Pg SEF cultured at 20℃, 28℃ and 37℃. The heatmap based on RNA-Seq results of the snake cells. The top 200 significant differentially expressed genes from the comparisons of 28℃ vs 20℃ and 28℃ vs 37℃ were selected, with the overlapping 80 genes shown in the heatmap.

**Supplemental Figure 4.**
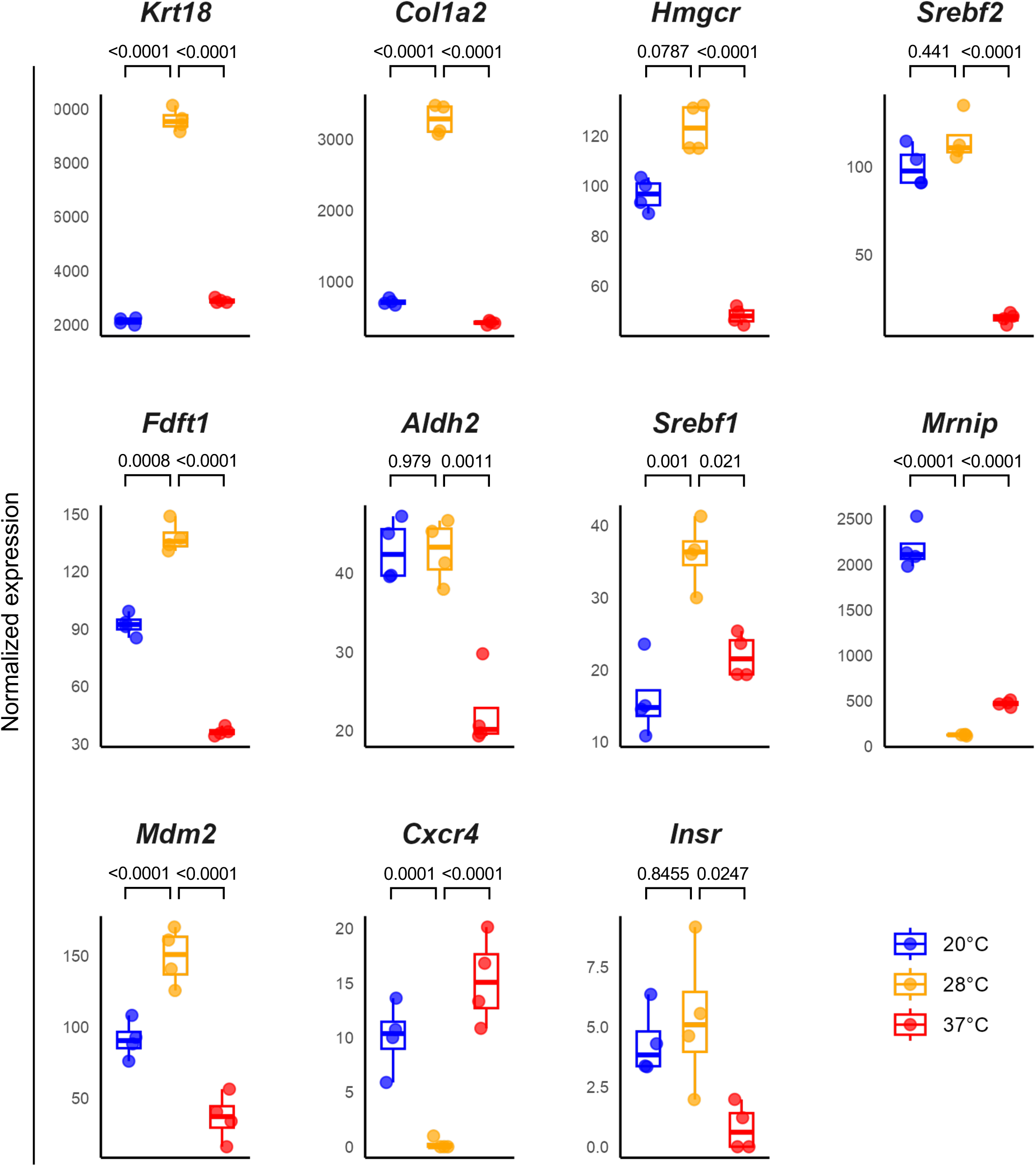
Differential expression of representative genes associated with enriched GO Biological Processes. Box plots showing the differential expression of genes related to cytoskeleton (*Krt18*), extracellular matrix (*Col1a2*), metabolism (*Hmgcr*, *Srebf2*, *Fdft1*, *Aldh2*, *Srebf1*), DNA repair (*Mrnip*, *Mdm2*), immune response (*Cxcr4*) and insulin receptor (*Insr*) signaling. Normalized expression and adjusted p-values are derived from DESeq2.

**Supplemental Figure 5.**
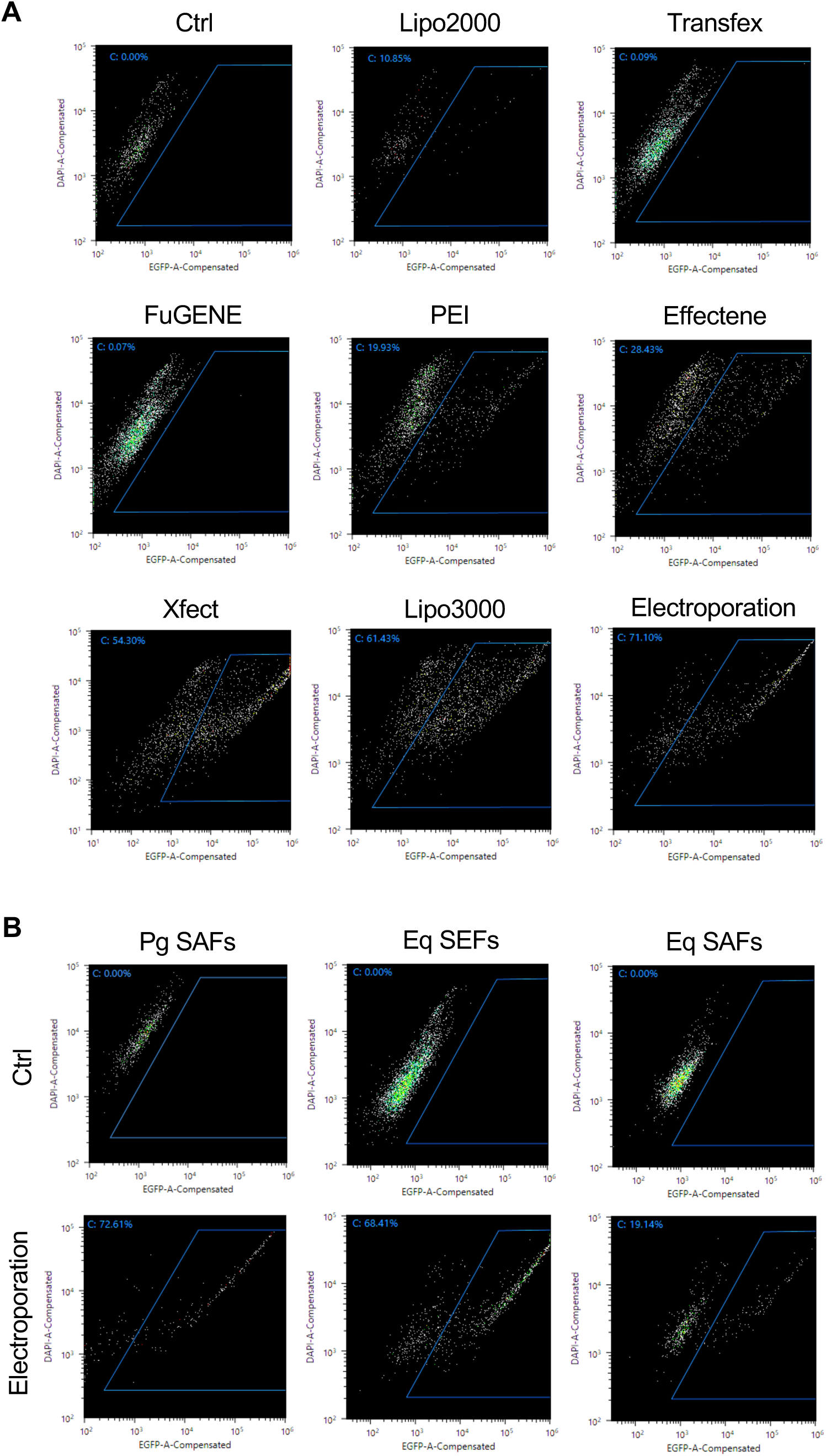
Representative results of the flow cytometry to validate the EGFP fluorescence, corresponding to Fig. 4. (A) The percentage of the EGFP-transfected Pg SEFs was validated by fluorescence-activated cell sorting (FACS). (B) The percentages of the EGFP-electroprated Pg SAFs, Eq SEFs and Eq SAFs were measured by FACS.

